# A high-quality Genome and Comparison of Short versus Long Read Transcriptome of the Palaearctic duck *Aythya fuligula* (Tufted Duck)

**DOI:** 10.1101/2021.02.24.432697

**Authors:** Ralf C Mueller, Patrik Ellström, Kerstin Howe, Marcela Uliano-Silva, Richard I Kuo, Katarzyna Miedzinska, Amanda Warr, Olivier Fedrigo, Bettina Haase, Jacquelyn Mountcastle, William Chow, James Torrance, Jonathan Wood, Josef D Järhult, Mahmoud M Naguib, Björn Olsen, Erich D Jarvis, Jacqueline Smith, Lél Eöry, Robert HS Kraus

## Abstract

**Background:** The tufted duck is a non-model organism that suffers high mortality in highly pathogenic avian influenza outbreaks. It belongs to the same bird family (Anatidae) as the mallard, one of the best-studied natural hosts of low-pathogenic avian influenza viruses. Studies in non-model bird species are crucial to disentangle the role of the host response in avian influenza virus infection in the natural reservoir. Such endeavour requires a high-quality genome assembly and transcriptome.

**Results:** This study presents the first high-quality, chromosome-level reference genome assembly of the tufted duck using the Vertebrate Genomes Project pipeline. We sequenced RNA (cDNA) from brain, ileum, lung, ovary, spleen and testis using Illumina short-read and PacBio long-read sequencing platforms, which was used for annotation. We found 34 autosomes plus Z and W sex chromosomes in the curated genome assembly, with 99.6% of the sequence assigned to chromosomes. Functional annotation revealed 14,099 protein-coding genes that generate 111,934 transcripts, which implies an average of 7.9 isoforms per gene. We also identified 246 small RNA families.

**Conclusions:** This annotated genome contributes to continuing research into the host response in avian influenza virus infections in a natural reservoir. Our findings from a comparison between short-read and long-read reference transcriptomics contribute to a deeper understanding of these competing options. In this study, both technologies complemented each other. We expect this annotation to be a foundation for further comparative and evolutionary genomic studies, including many waterfowl relatives with differing susceptibilities to the avian influenza virus.

## Background

The tufted duck (*Aythya fuligula*) is a non-model organism that has received attention because of its role in the zoonotic ecology of avian influenza A viruses (AIV). As a member of the Anatidae family of ducks, it is closely related to the mallard (*Anas platyrhynchos*), the primary natural host of AIV [1–5]. However, in contrast to mallards, which carry this virus largely asymptomatically, tufted ducks are less commonly infected but suffer high mortality in outbreaks of highly pathogenic avian influenza (HPAI) [6–9]. The tufted duck is a diving duck. Its breeding range is throughout northern Eurasia where it is largely a seasonal short-distance migrant. Although it generally feeds in deeper waters than mallards and other dabbling ducks, it generally shares its habitat and roosting areas with these and many other waterfowl species, even leading to a high rate of interspecific hybridisation [10, 11]. Hence, given a frequent exposure to AIVs circulating in such habitats, the differences in susceptibility to-and outcome of AIV infection between these species are likely related to genetic differences affecting, e.g., receptor expression or host response. For example, the resistance of mallards against severe HPAI infection has been partially attributed to the presence of the RIG-I gene and its strong interferon response, in contrast to chickens that lack this gene and develop severe disease upon HPAI infection [12]. Future studies in non-model bird species such as the tufted duck are important to disentangle the role of the host response and other genetic factors in AIV infection and aid in our understanding of the zoonotic ecology of AIV in the natural reservoir. A prerequisite for such studies is a well-assembled and annotated genome and transcriptome [13].

Developments in omic sequencing technologies over the last two decades have revolutionised biology. Instead of studying single genes and their products, whole genomes and transcriptomes can now be readily sequenced and assembled at a lower cost than before. Massive parallelisation and high throughput in next-generation sequencing (NGS) have decreased sequencing costs and ultimately increased sequencing depth [14]. This allows for whole-genome sequencing of any species and opens up new possibilities for in-depth studies related to infection biology and host response to external stressors beyond model species in which a rich genetic toolbox can be deployed [15, 16]. Non-model species are frequently understudied, yet they are exposed to environmental stressors like infectious diseases, which they can transmit to livestock and humans [17]. Approximately 70% of human infectious diseases are zoonoses [18, 19]. An in-depth understanding of a pathogen’s zoonotic ecology in the animal reservoir is important to prevent human infections. This, in turn, requires studies of host-pathogen interactions at the genetic level. NGS helps bridge the gap between model and non-model species [20] and with third-generation long-read based sequencing, high-quality reference genomes are now also available for non-model organisms, as in the Vertebrate Genomes Project (VGP) [21]. This is supplemented by affordable long-read RNA sequencing, making *de novo* assembled transcriptomes unnecessary [22].

In *de novo* sequencing/assembly, high sequencing depth with short reads (< 250 bp) is used in an attempt to recover whole genomes, which decreases uncertainty during the assembly process. While this results in the possibility of studying previously uncharacterised species without relying on reference genomes, there are still limitations. For instance, short reads make it difficult to assemble and identify very similar genes (e.g., paralogs), regions of low complexity (e.g., tandem repeats), or GC-rich microchromosomes of birds [16, 23]. Illumina sequencing produces short reads, which often fail to assemble these difficult regions leading to artefacts, misassemblies or incorrect sequence reconstructions [21, 24, 25]. On the other hand, the high sequencing base call quality accurately recovers nucleotide sequences in critical regions, e.g., splice junction sites. The introduction of PacBio’s Single Molecule, Real-Time (SMRT) sequencing offers the opportunity to sequence very long reads (N50 > 10kb) of unfragmented DNA [26, 27] leading to lower assembly structural errors and ambiguity. As there is no prior amplification needed, there is also no PCR-induced variance of coverage in the sequences. However, error rates can be high for continuous long-read (CLR) technologies, resulting in more singlenucleotide errors. This can be corrected by increasing sequencing depth, using specialised polishing algorithms with long reads or high accuracy short reads [21]. Exploiting each technology’s strength facilitates more in-depth and unbiased genome studies while also enabling us to draw much more robust and detailed conclusions regarding research questions, e.g., identifying genes and gene products that previously escaped detection [24].

Transcriptome annotation of the gnome used to be constrained to either low-throughput and costly cDNA cloning or Illumina’s high-throughput short-read RNA sequencing (RNA-Seq). High-quality short-read RNA sequencing combined with a reference genome allows for a reasonable transcriptome reconstruction. However, there are some caveats: Due to alternative splicing, a single gene can have multiple alternative variants (isoforms) and as a consequence can be translated into proteins with different functions [25, 28]. Illumina short reads need to be assembled into transcripts, which can lead to incompleteness and errors in transcript model reconstruction. This biases the correct inference of isoforms and thus misses the transcriptome’s complexity [25, 29]. Furthermore, since short-read sequencing is also limited in GC-rich regions and regions of low sequence complexity [24], not all transcripts are recovered. In contrast, in full-length transcript isoform sequencing (e.g., PacBio Iso-Seq) of mRNA, the single direct reads from 5’ to 3’ usually do not need to be assembled and, thus prevent assembly ambiguities, which facilitates the detection of novel isoforms [30, 31] and accurate reconstruction of transcript structure [24]. This third-generation transcript sequencing technology also allows for a much more detailed functional annotation. It is crucial to recover as many isoforms as possible for functional studies [32–34]. For example, the immunome comprises a set of genes associated with immune processes and is a heavily complex portion of the genome [22, 35]. Thus, the benefits of 3^rd^ generation sequencing must be exploited to their fullest potential to facilitate studies in bird immunogenomics [36] and improve databases such as the Avian Immunome DB for comparative analyses in non-model bird species [37].

## Data Description

This study presents the first high-quality, chromosome-level reference genome assembly of the tufted duck using the VGP pipeline [21], which we annotated with short- and long-read transcripts from six different tissues, and short reads from small RNAs from the same tissues. We further contrasted different sequencing technologies and bioinformatics pipelines, and tissue-specific comparisons of expressed genes. This annotation serves as a foundation for further comparative and evolutionary genomics, and gene expression experiments in the tufted duck and its many waterfowl relatives with different susceptibilities to AIV [36, 38].

## Analyses

### Reference genome assembly

The reference genome fully met VGP’s 6.7.P2.Q40.C99 metric standards [21]. This includes contig NG50 > 10^6^ bp, scaffold NG50 > 10^7^ bp, most of the genome separated into haplotypes (P2), Phred-scaled base accuracy > Q40, and 99.5% of the assembly assigned to chromosomes. The contig NG50 of the tufted duck genome assembly was 17.8 Mb. The curation resulted in 34 removals of misjoins, 34 joins previously missed, and four removals of false duplications. This reduced the primary assembly length by 0.8% and increased the scaffold NG50 by 18.7% to 85.9 Mb whilst decreasing the scaffold number from 123 to 105. Chromosomal-scale units were identified and named by size. Out of the expected 39 chromosome pairs according to the karyotype [39], 34 autosomes plus Z and W could be identified, with 99.6% of the sequence assigned to them (Tab. 1, Fig. 1).

**Fig. 1.**
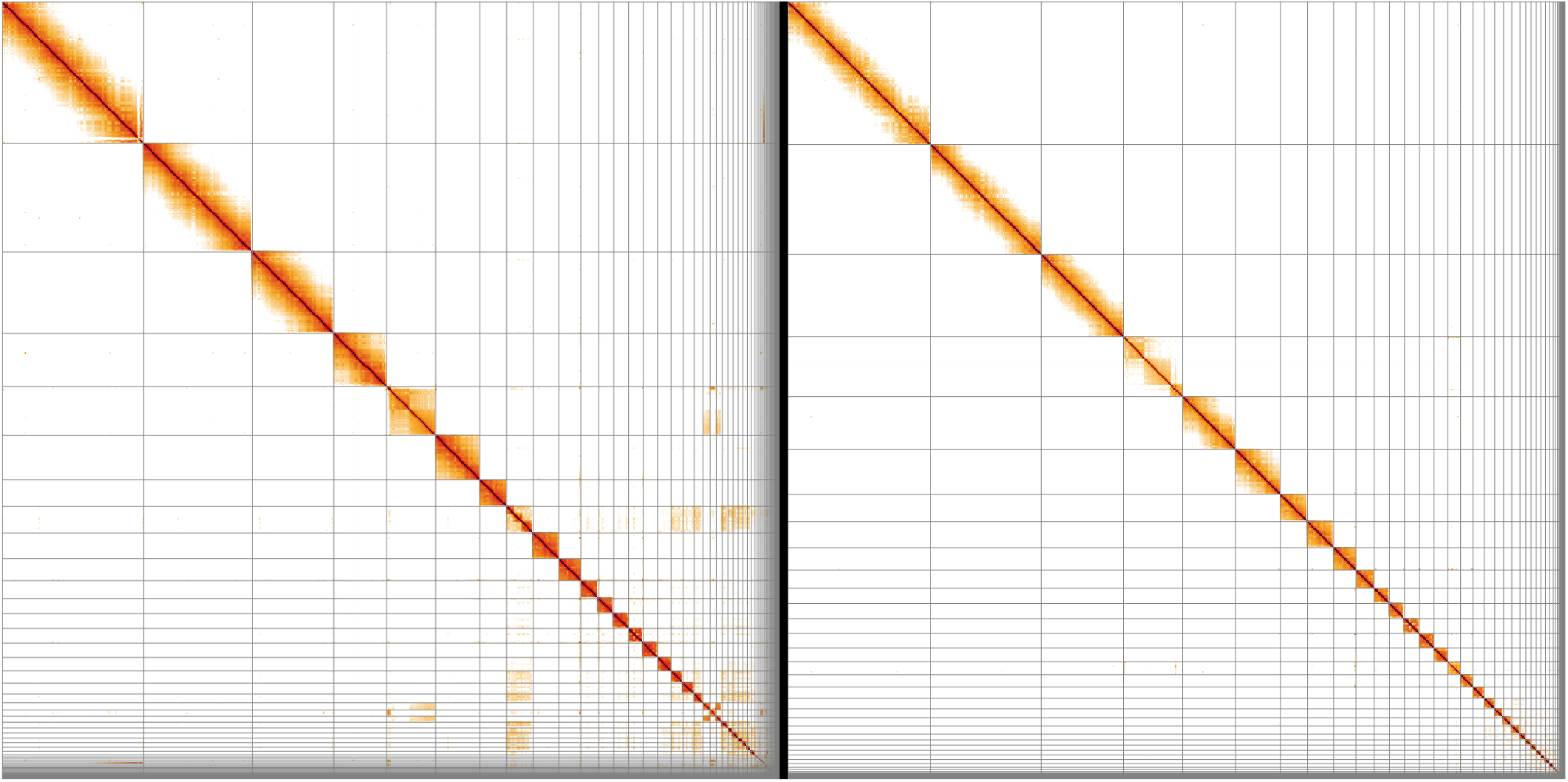
HiGlass Hi-C 2D maps of the tufted duck genome assembly before (left) and after (right) manual curation. Off-diagonal hits indicate missing joins, which have been corrected during curation. Broken patterns within scaffolds (e.g., at the end of the first scaffold pre-curation) can indicate intra-scaffold misassemblies, which were also addressed during curation. They can, however, also be features of the respective chromosome, as in the 4^th^ post-curation scaffold the structure of which was corrected and asserted during curation.

**Tab. 1.** Assembly statistics of the tufted duck genome.

|  |  |
| --- | --- |
| Genome coverage [x] | 64.03 |
| Total sequence length [bp] | 1,127,004,725 |
| Ungapped sequence length [bp] | 1,117,587,328 |
| Number of scaffolds | 105 |
| Scaffolds assigned to chromosomes | 36 + 1 mitochondrion |
| Unplaced scaffolds | 68 |
| Contig NG50 [bp] | 17,816,505 |
| Scaffold NG50 [bp] | 85,905,639 |
| Base-call error in 10 kb | less than 1 nucleotide |

The raw data has been deposited in the GenomeArk repository [40]. The curated primary assembly has been deposited in NCBI under accession number GCF_009819795.1 [41] and can be browsed in the Genome Data Viewer [42].

### Gene/transcript annotation with Illumina RNA-Seq and PacBio Iso-Seq reads

With the Illumina RNA-Seq, an average of 97.65% (SD = 0.51%) of the reads were retained after adapter and quality trimming. For the PacBio Iso-Seq, an average of 80.57% (SD = 3.84%) full-length non-chimeric reads were retained after error correction (Tab. 2).

On average, HISAT2 could map 93.21% (SD = 0.64%) Illumina RNA-Seq short reads and Minimap2 could map 97.39% (SD = 1.34%) PacBio Iso-Seq long reads to the reference genome. The average read length of the long-read data was 1214.0 nt (SD = 262.0), an order of magnitude longer than that of the short reads (average 131.5 nt, SD = 0.7). However, the average number of reads was almost 600-fold higher with short-read data (129,411,992; SD = 11,989,532) compared to long-read data (217,293; SD = 143,219), implying a 60 times higher sequencing depth with the short reads. Not surprisingly, StringTie2 (in the short-read pipeline) assembled more transcript models and inferred more genes and exons than the long-read pipeline. This was true for all tissues except in the lung; more transcript models were inferred in the long-read pipeline. Lung RNA (cDNA) was sequenced on two Zero-mode waveguides (ZMW) and produced the highest numbers of subreads and full-length, non-chimeric reads (FLNC) after processing. The highest number of genes in each pipeline was predicted for ovary (Illumina) and brain (PacBio). The highest number of transcript models was predicted for ovary (Illumina) and lung (PacBio). The same pattern applies to predicted exons (Tab. 3).

In the Illumina short-read RNA-Seq annotations, more exons per gene were recovered on average than in the long-read pipeline (Tab. 3). The distribution of exons per gene in both pipelines generally followed the same pattern with a decreasing number of genes as the exons per gene increased. However, there is a remarkable difference in the short-read data, which recovered more two-exon genes than single-exon genes for all tissues (Fig. 2).

**Fig. 2.**
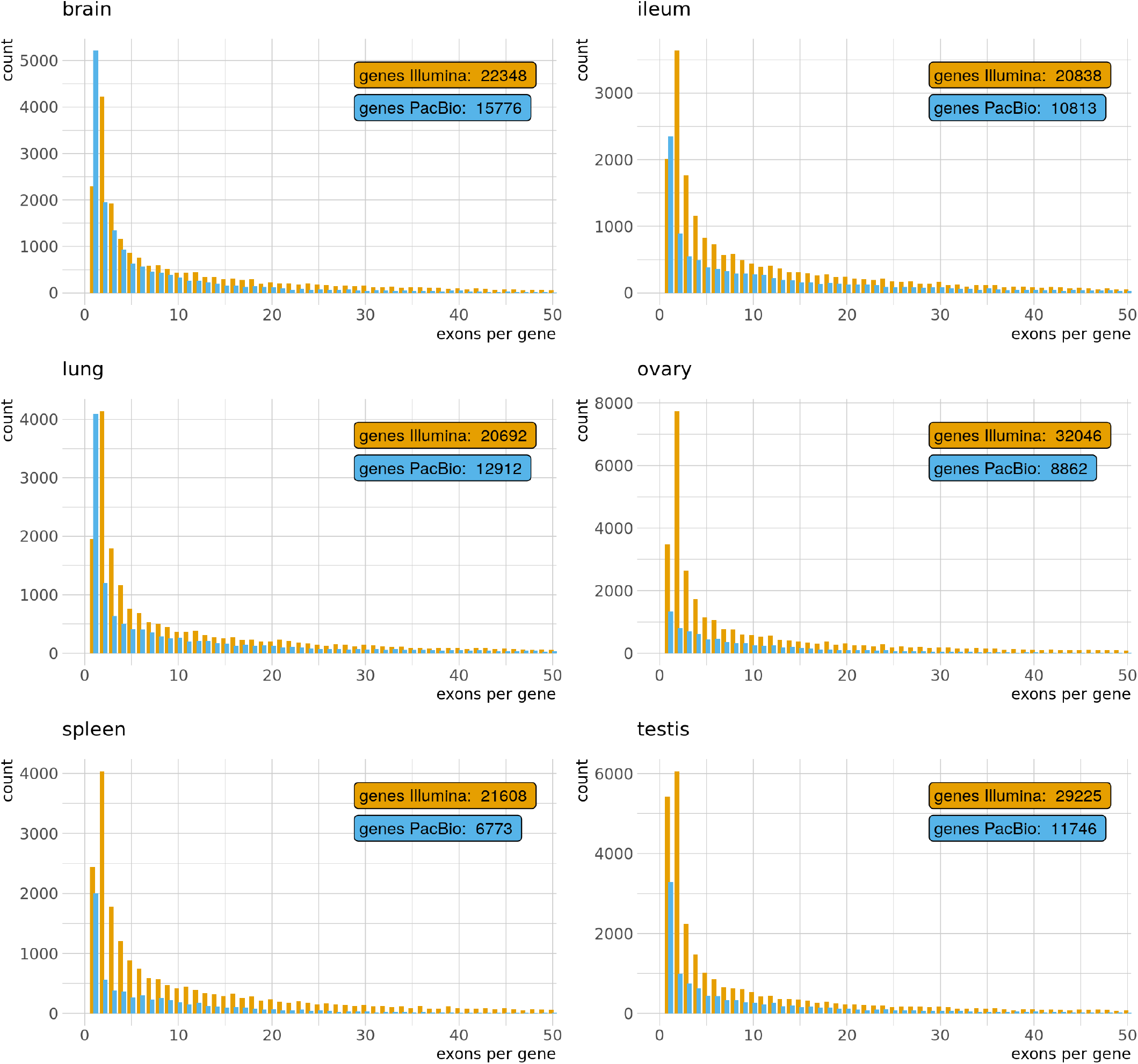
Distribution of single-and multi-exon genes per tissue and pipeline. Only the first 50 groups are shown.

**Tab. 2.** Illumina RNA-Seq reads before and after trimming, and PacBio Iso-Seq reads before and after error correction. Zero-mode waveguide (ZMW) yield and the number of full-length, non-chimeric reads (FLNC). Two ZMW were used for lung and ovary.

| Platform |  | brain | ileum | lung | ovary | spleen | testis |
| --- | --- | --- | --- | --- | --- | --- | --- |
| Illumina | Number of raw reads (PE) | 71,648,303 | 72,410,462 | 63,191,981 | 72,442,392 | 68,315,029 | 53,747,262 |
|  | Number of trimmed (paired) | 70,025,835 | 70,902,250 | 61,059,701 | 70,892,177 | 66,813,691 | 52,652,923 |
| PacBio | Number of subreads | 9,249,099 | 12,363,369 | 19,206,097 | 1,279,561 | 8,401,115 | 12,616,953 |
|  | Number of ccs | 158,698 | 415,314 | 529,108 | 68,112 | 167,077 | 288,984 |
|  | ZMW yield [%] | 15.87 | 41.53 | 26.46 | 3.41 | 16.71 | 28.90 |
|  | Average number of passes | 58.3 | 29.8 | 36.3 | 18.8 | 50.3 | 43.7 |
|  | Number of FLNC | 133,684 | 343,634 | 432,817 | 49,887 | 134,124 | 234,423 |

In the PacBio long-read Iso-Seq annotations, more transcripts per gene were recovered on average than in the short-read pipeline (Tab. 3). The distribution of transcripts per gene followed the same pattern with a decreasing number of genes as the transcripts per gene increased. However, these numbers had a slower decay in the long-read data, indicating a better recovery of multi-transcript genes (Fig. 3).

**Fig. 3.**
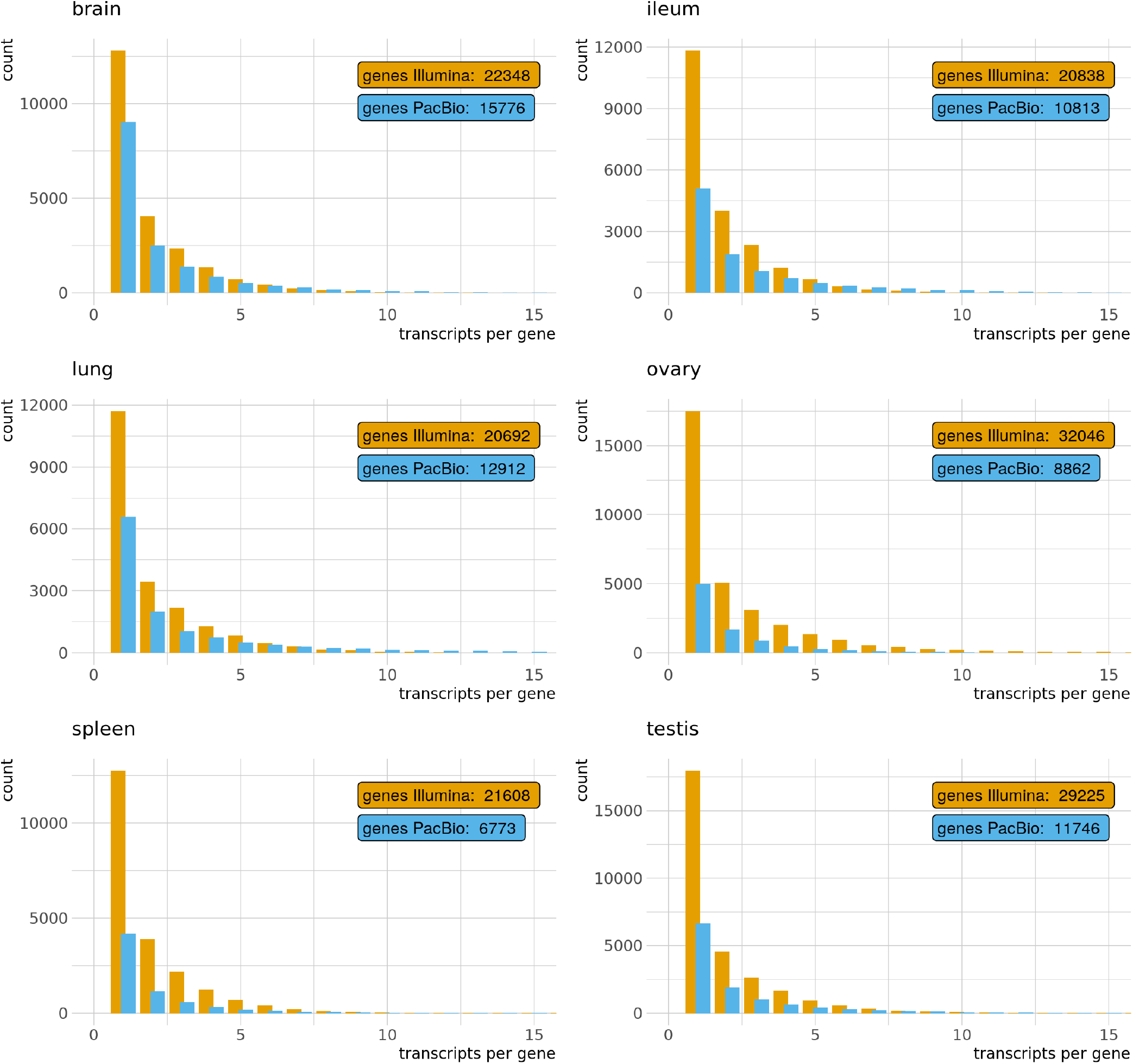
Distribution of single-and multi-transcript genes per tissue and pipeline. Only the first 15 groups are shown.

**Tab. 3.** Annotation per tissue and pipeline.

|  |  | brain | ileum | lung | ovary | spleen | testis |
| --- | --- | --- | --- | --- | --- | --- | --- |
| Mapped [%] | Illumina | 92.92 | 92.98 | 93.42 | 94.33 | 92.44 | 93.19 |
|  | PacBio | 96.66 | 98.13 | 97.92 | 99.40 | 95.67 | 96.57 |
| Number of genes | Illumina | 22,348 | 20,838 | 20,692 | 32,046 | 21,608 | 29,225 |
|  | PacBio | 15,776 | 10,813 | 12,912 | 8,862 | 6,773 | 11,746 |
| Number of transcripts | Illumina | 44,808 | 40,968 | 43,877 | 77,997 | 42,000 | 57,758 |
|  | PacBio | 37,601 | 35,284 | 46,587 | 19,513 | 14,030 | 28,852 |
| Number of exons | Illumina | 422,741 | 395,719 | 412,225 | 569,566 | 383,679 | 483,444 |
|  | PacBio | 138,038 | 217,391 | 243,566 | 119,080 | 71,211 | 160,380 |
| Exons per gene (average) | Illumina | 18.9 | 19.0 | 19.9 | 17.8 | 17.8 | 16.5 |
|  | PacBio | 8.7 | 20.1 | 18.9 | 13.4 | 10.5 | 13.7 |
| Transcripts per gene (average) | Illumina | 2.0 | 2.0 | 2.1 | 2.4 | 1.9 | 2.0 |
|  | PacBio | 2.4 | 3.3 | 3.6 | 2.2 | 2.1 | 2.5 |

### Functional annotation of merged transcripts

After merging all twelve annotations (six tissues with short reads and long reads), 345,870 transcripts with 2,381,662 exons were predicted in 49,746 genes (Tab. 4). Of these, 178,198 transcripts (or 16,758 genes) were predicted by CPC2 to be protein-coding. UniProt hits were found for 208,274 transcripts (or 17,911 protein-coding genes). Of these, 4,766 genes had no long-read support and 432 genes no short-read support in the data. The number of protein-coding genes predicted by CPC2 and number of hits in UniProt overlapped for 15,103 genes (Fig. S1). Conservative filtering of the annotation (ignoring features flagged as nonsense-mediated decay and only keeping full-length hits in UniRef50 which matched with at least 90%) resulted in 111,934 transcripts and 14,099 protein-coding genes. This result implies an average of 7.9 isoforms per gene (Tab. 4).

**Tab. 4.**
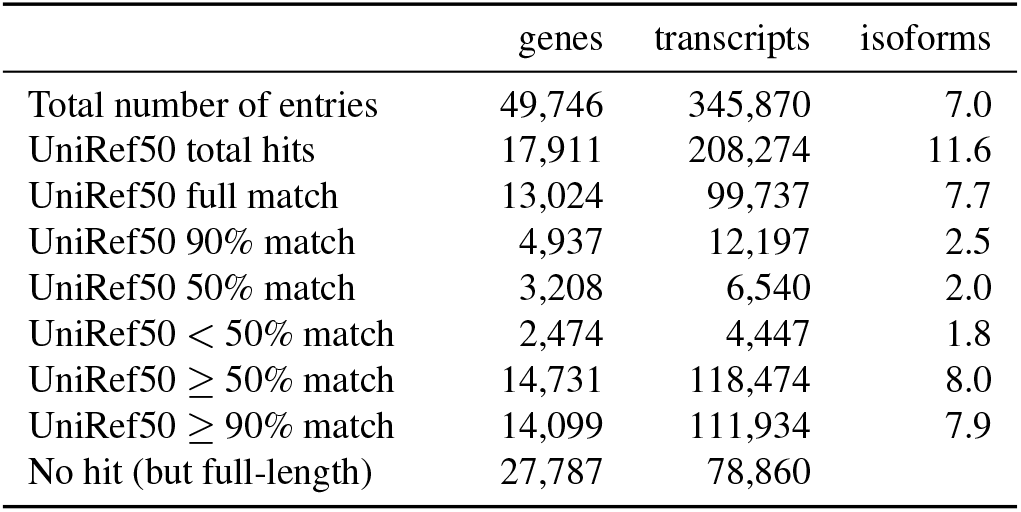
Functional annotation categorised by different matches. Note: UniRef50 total hits also includes 5’ degraded records, whereas the match classes only include full-length records.

|  | genes | transcripts | isoforms |
| --- | --- | --- | --- |
| Total number of entries | 49,746 | 345,870 | 7.0 |
| UniRef50 total hits | 17,911 | 208,274 | 11.6 |
| UniRef50 full match | 13,024 | 99,737 | 7.7 |
| UniRef50 90% match | 4,937 | 12,197 | 2.5 |
| UniRef50 50% match | 3,208 | 6,540 | 2.0 |
| UniRef50 < 50% match | 2,474 | 4,447 | 1.8 |
| UniRef50 ≥ 50% match | 14,731 | 118,474 | 8.0 |
| UniRef50 ≥ 90% match | 14,099 | 111,934 | 7.9 |
| No hit (but full-length) | 27,787 | 78,860 |  |

Out of 78,860 full-length transcripts with no UniRef50 hit, 62,147 were flagged as potentially protein-coding (and the remainder as nonsense-mediated decay), and 26,489 as single-exonic while the remainder as composed of two or more exons.

### Tissue-specific expression and intersections

The number of supported protein-coding genes was generally higher with Illumina transcriptome reconstruction than with that of PacBio. The short-read pipeline supported 11,165 annotated genes across all tissues. The highest number of exclusively expressed genes was in testis (988) followed by ovary, brain, spleen, ileum and lung (Fig. 4). The long-read pipeline supported 2,475 annotated genes across all tissues. The highest number of exclusively expressed genes was in brain (779) followed by testis, ileum, lung, ovary and spleen (Fig. 5). Overlap of expressed genes from each pipeline can be found in Fig. S2.

**Fig. 4.**
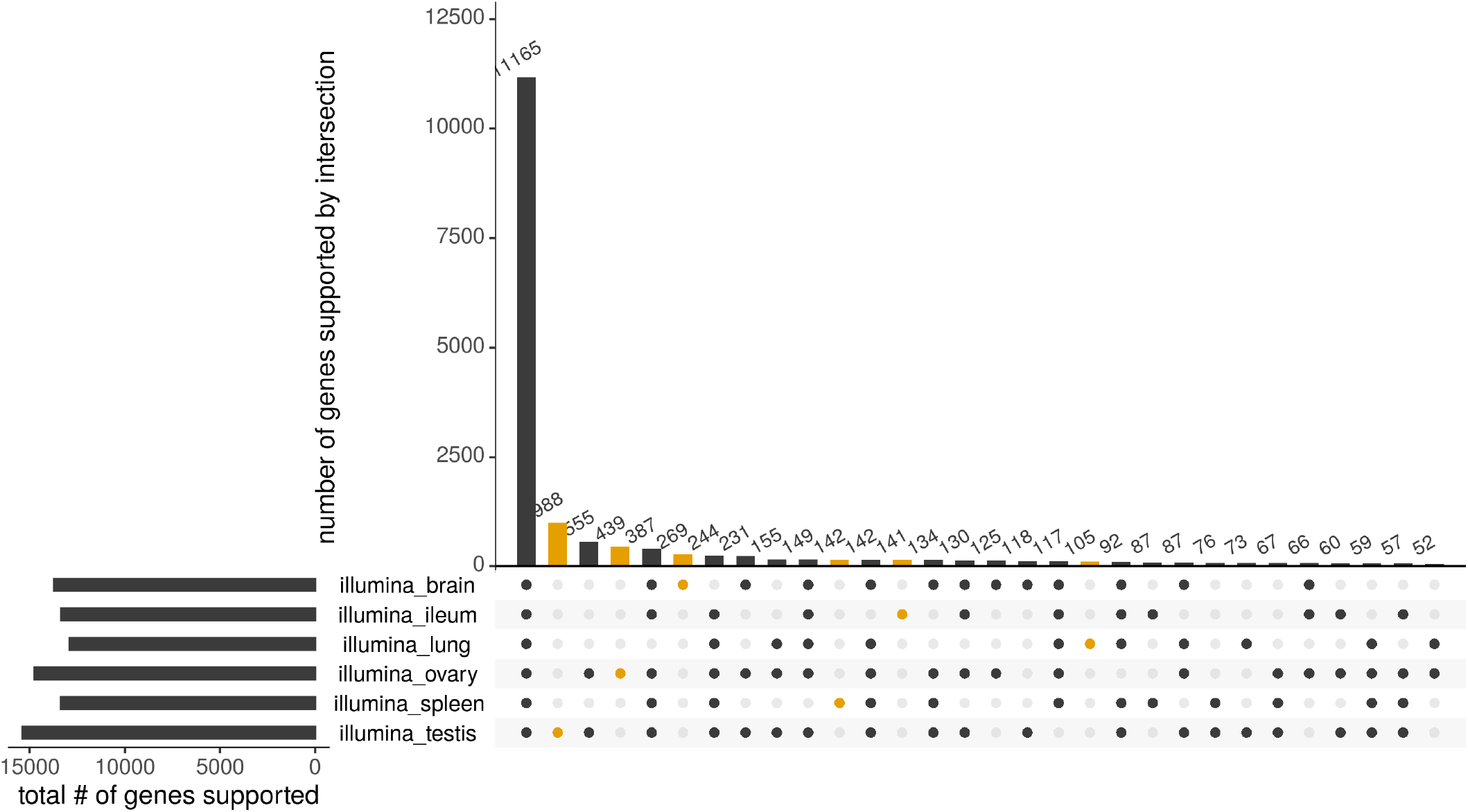
In the short-read data set, the highest total number of supported genes was found in testes (left panel, bottom), followed by ovary, brain, spleen, ileum and lung. All six tissues intersected in 11,165 genes (main panel, left). The highest number of exclusively supported genes was also found in testes (988), and followed the same order as the total number of genes (main panel, yellow).

**Fig. 5.**
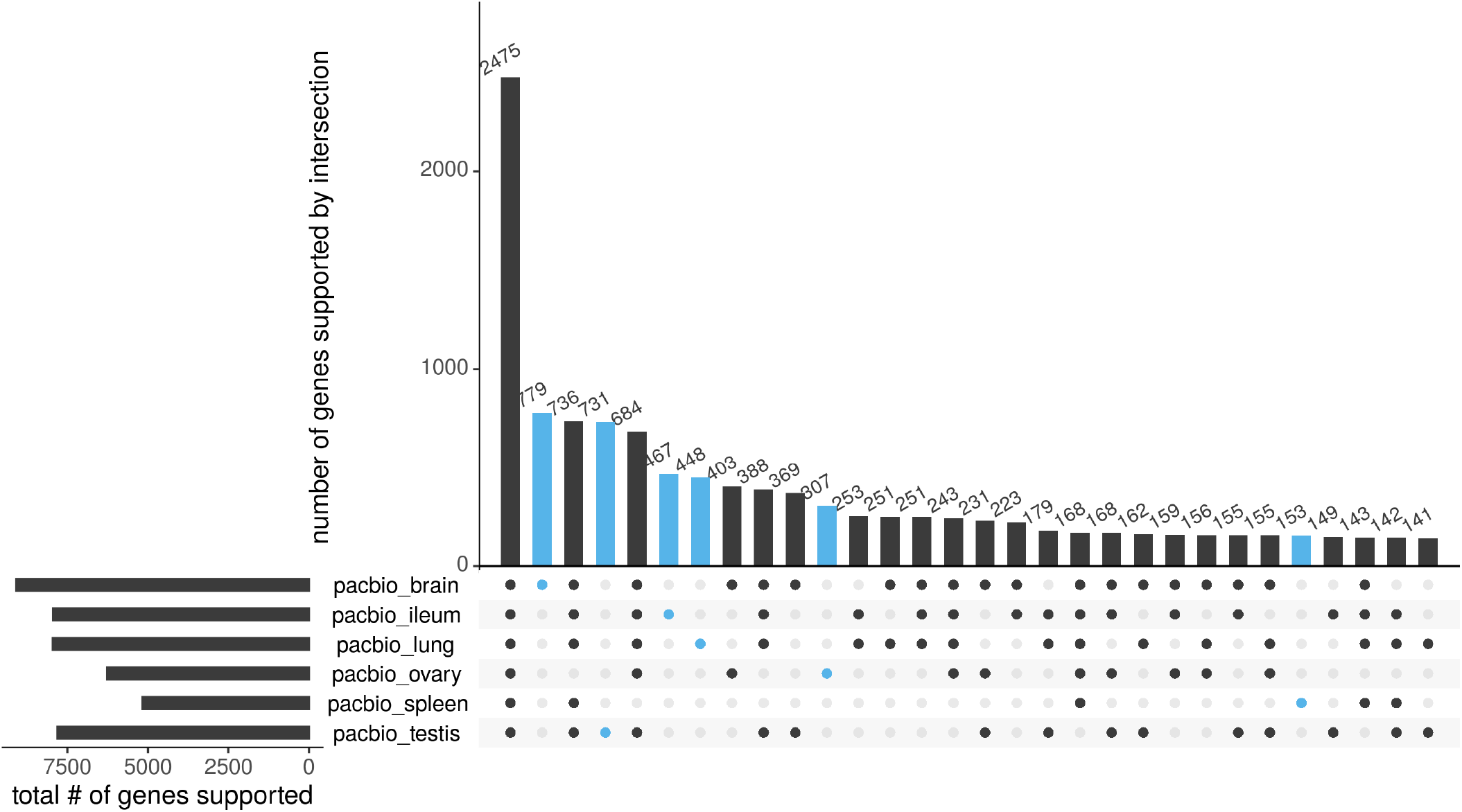
In the long-read data set, the highest total number of supported genes was found in brain (left panel, top), followed by lung, ileum, testes, ovary and spleen. All six tissues intersected in 2,475 genes (main panel, left). The highest number of exclusively supported genes was found in brain (779), followed by testes, ileum, lung, ovary and spleen (main panel, blue).

### Small RNA analyses

For the TruSeq small RNA sequencing data, 95.81% (SD = 3.51%) of the reads were retained (on average) after adapter and quality trimming (Tab. 5). Overall, STAR could map on average 93.91% (SD = 5.28%) of these reads to the reference genome, which divides into 67.99% (SD = 13.75%) uniquely mapped reads and 25.92% (SD = 12.08%) multi-mapped reads (up to 10 loci). Cufflinks predicted the highest number of genes in the spleen (33,133) followed by testis (31,205). The remaining tissues had much lower numbers of genes ranging from 8,441 (ileum) to 13,606 (brain). The same pattern applies to the number of predicted transcripts and exons (Tab. 5).

**Tab. 5.**
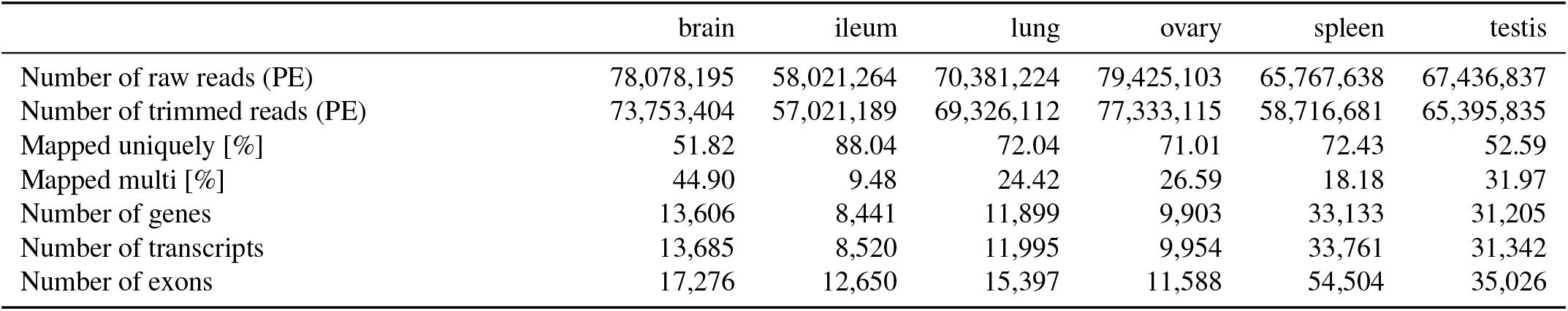
Small RNA read processing and assembly stats.

|  | brain | ileum | lung | ovary | spleen | testis |
| --- | --- | --- | --- | --- | --- | --- |
| Number of raw reads (PE) | 78,078,195 | 58,021,264 | 70,381,224 | 79,425,103 | 65,767,638 | 67,436,837 |
| Number of trimmed reads (PE) | 73,753,404 | 57,021,189 | 69,326,112 | 77,333,115 | 58,716,681 | 65,395,835 |
| Mapped uniquely [%] | 51.82 | 88.04 | 72.04 | 71.01 | 72.43 | 52.59 |
| Mapped multi [%] | 44.90 | 9.48 | 24.42 | 26.59 | 18.18 | 31.97 |
| Number of genes | 13,606 | 8,441 | 11,899 | 9,903 | 33,133 | 31,205 |
| Number of transcripts | 13,685 | 8,520 | 11,995 | 9,954 | 33,761 | 31,342 |
| Number of exons | 17,276 | 12,650 | 15,397 | 11,588 | 54,504 | 35,026 |

On average, each transcript was composed of 1.3 exons (SD = 0.2 exons). The distribution of single-exon and multiexon transcripts shows a clear trend towards single-exon transcripts and a quickly diminishing number of multi-exon transcripts. However, this was less pronounced in spleen but even more so in testis (Fig. 6). The generally decaying length distribution of transcripts shows two clear peaks at 20 bp and 50 bp except for testis with the first peak at 30 bp. Except for spleen and testis data, there are very few transcripts > 200 bp (Fig. S3).

**Fig. 6.**
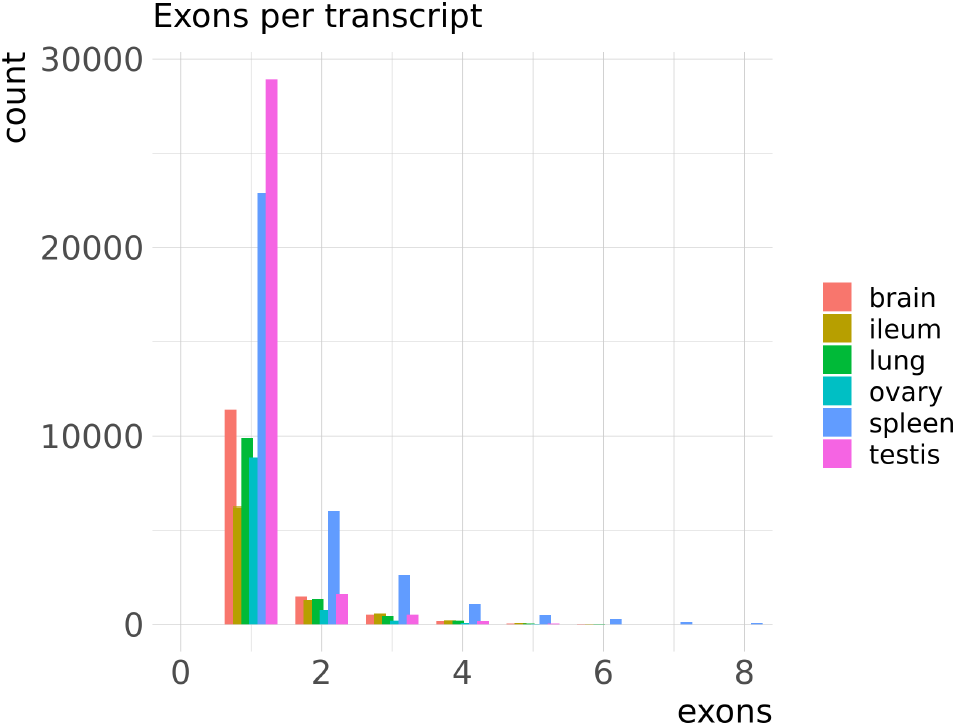
Distribution of single-exon and multi-exon small RNA transcripts for each tissue.

**Table 6.** Results of cmscan on assembled small RNA transcripts after filtering. Intersection refers to small RNAs predicted by the *in silico* genome scan. Additional refers to annotated small RNAs which were not detected by CMSCAN in the reference genome.

|  | brain | ileum | lung | ovary | spleen | testis |
| --- | --- | --- | --- | --- | --- | --- |
| *in vitro* | 328 | 294 | 317 | 312 | 345 | 369 |
| intersection | 310 | 274 | 295 | 293 | 315 | 345 |
| additional | 18 | 20 | 22 | 19 | 30 | 24 |

Scanning the genome *(in silico)* for Rfam’s covariance models of RNAs resulted in 1,234 hits. The same scan on the assembled small RNA transcripts *(in vitro)* revealed an average of 346.5 hits (SD = 26.9) across all tissues. After removing lower-scoring overlaps and hits with E-value larger than 5.0E-4 from the cmscan result, 1,076 distinct features were predicted in the tufted duck genome. In the tissue’s small RNA assemblies, 327.5 (SD = 26.5) features were annotated, on average, with the same filtering (Tab. 6). On average 93.25% (SD = 1.09%) of the *in vitro* annotated features were predicted by the *in silico* scan. Furthermore, 22.2 (SD = 2.3) features were annotated on average, which were not predicted *in silico* (Tab. 6).

After further filtering for unique RNA families (Rfam accession numbers/covariance models), 306 distinct RNA families were predicted in the genome, with 246 annotated in all tissues (pooled). The number of predicted and annotated covariance models overlapped for 237 RNA families, while 69 were only identified in the genome scan, and nine were only identified in the pooled tissue annotations.

## Discussion

In this study, we present the first chromosome-level reference genome assembly of the tufted duck. The genome contiguity is on par with other reference bird genome assemblies that used long reads like mallard [43, 44], chicken [45], and recent VGP-pipeline generated zebra finch [46]. The assembly’s contig NG50 of 17.8 Mbp is comparable to chicken (17.7 Mbp) but considerably higher than in mallard (5.7 Mbp) and zebra finch (4.4 Mbp). The assembly’s scaffold NG50 of 85.9 Mbp is higher than in mallard (76.3 Mbp) and zebra finch (70.9 Mbp), and considerably higher than in chicken (20.8 Mbp). All our mapping results from the Illumina short-read and PacBio long-read RNA pipelines confirm full-adherence to the VGP 6.7.P2.Q40.C99 standard, which also implies transcript mappability > 80% [21].

The higher numbers of recovered genes, transcripts and exons from the short-read transcript pipeline may be mainly explained by the higher sequencing depth and further reinforced by the different RNA preparation protocol. With Illumina, virtually all trimmed, paired-end reads were kept for mapping to the genome while with PacBio, only 5’ cap-selected and full-length, non-chimeric reads were kept. In the short-read pipeline, more two-exon genes than single-exon genes were predicted for all tissues, and it appears as if some transcripts could not be assembled entirely or StringTie2 tried to “avoid” single-exon genes. Transcript model reconstruction in StringTie2 is based on the concept of extending short reads to create so-called super-reads [47], which seems appropriate for whole-genome assemblies. In transcriptomics, however, multiple splice variants are possible, and with the super-read concept in mind, it may thus be possible that StringTie2 discards a single-exon transcript model in favour of an alternative multi-exon splice variant containing the same exon. This, in turn, might have a substantial impact on the functional annotation based on transcript models solely assembled with short reads. Real single-exon transcripts might be missed, or even worse, false-positive multiexon transcript models might be reconstructed. We merged transcript models of both pipelines with higher priority on transcript end sites for the long-read-inferred transcript models to mitigate this effect. The average number of transcripts in the long-read pipeline often almost matched or even exceeded (lung) the average number of transcripts in the shortread pipeline. Furthermore, the long-read pipeline recovered more transcripts per gene (isoforms) on average. Taken together, this is remarkable considering the 60-fold higher sequencing depth in the short-read pipeline. It also corroborates the strength of PacBio Iso-Seq, which seems to better reflect the complexity of the transcriptome, considering that transcripts did not need to be assembled but were sequenced full-length.

While both the number and quality of published vertebrate genome assemblies are rising, hardly any are complemented by a transcriptome of multiple tissues from the same individual [25, 48]. Automated annotations (e.g., the NCBI Eukaryotic Genome Annotation Pipeline, [49]) provide reasonable *in silico* predictions of coding potential; however, RNA (cDNA) sequencing adds evidence for expressed genes. Based on the inferred transcripts in this study, a total of 14,099 protein-coding genes could be identified in the UniRef50 database after conservative filtering (at least 90% match), which is comparable to NCBI’s *in silico* prediction of 15,578 protein-coding genes [50]. The number of identified protein-coding genes in tufted duck is also in line with the prediction in other bird species like mallard (16,836; [51]), chicken (17,477; [52]) or zebra finch (16,197; [53]). CPC2 predicted 84.2% of the potentially protein-coding genes found in UniRef50, which would serve as a conservative estimate of the protein-coding potential by just looking at the reference genome. However, beyond the 17,911 genes annotated by UniRef, the annotation contains an additional 27,787 genes with protein-coding potential according to the TAMA ORF/NMD prediction pipeline, with these being potential candidates for further analyses. Gene and transcript identification in non-model species relies on annotations of preferably closely-related model organisms. However, protein-coding genes are mainly described by a single transcript and predominantly built on short-read and comparative data [25].

Besides a high-quality transcriptome for the tufted duck, this study also provides a tissue-specific expression atlas. In the short-read pipeline, there is a large drop in numbers of transcripts supported in all tissues to transcripts exclusively expressed in a single tissue or a few tissues. This distribution is much more balanced in the long-read pipeline and may indicate a better recovery of isoforms or a higher number of falsepositive assembled transcripts in the short-read pipeline.

The number of 306 Rfam small RNA families predicted (with 246 annotated) for the tufted duck in this study is comparable to 352 families predicted in chicken (Gallus gallus, version 5) in Rfam 14.4 [54]. The peaks of transcript length at 20 bp and 50 bp are in the area of microRNA (miRNA), which are usually 18 - 23 nt [55–57], and pre-miRNA which are in the range of 55 - 70 nt. The substantial variance across tissues in our data (many more genes, transcripts and exons in spleen and testis than in the other tissues) might be explained by Illumina sequencing bias introduced at the adapter-ligation step of cDNA library constructions [58, 59]. Furthermore, according to Illumina’s TruSeq Small RNA library preparation protocol, small RNA populations can vary significantly between different tissue types and species, and types and coverage vary depending on which bands are selected during gel excision. This is additional support of a strategy to sequence multiple tissues to obtain a fuller picture of small RNA expression in an organism. Gene duplications might explain the significant difference between unique mappings and multi mappings of small RNA to the genome across tissues. The *in silico* scan of small RNA in the genome could predict almost all small RNAs in the assembled transcripts. More importantly, however, 22.2 additional small RNAs were discovered on average in the *in vitro* scan which would otherwise have been unnoticed. Small (non-coding) RNA play an essential role in gene regulation, translation and chromosome structure [60, 61], and are often associated with diseases [62, 63]. This vital fraction of the genome is rarely validated *in vitro* in the genome and transcriptome annotation literature. However, small RNA studies have been continuously rising over the last 20 years from 1,966 publications in 1999 to 8,034 in 2019 (searching for “small RNA” on Web of Science [64]). Relying on *in silico* prediction of small RNA alone can lead to misinterpretation of pathways and gene regulation, and sequencing small RNA of non-model organisms is, therefore, an advancement for genome annotation [65]. We strongly encourage *in vitro* sequencing of small RNA in *de novo* genome and transcriptome studies, or otherwise, the scientific community will miss such detail that will be important to decipher relevant differences in the biology between species.

### Potential implications

This study presents the first high-quality reference genome assembly of the non-model tufted duck species. It is complemented by coding and small non-coding RNA transcriptome annotation from six different tissues. The genome assembly contributes to the VGP’s ongoing mission to generate near error-free and complete genome assemblies of all extant vertebrate species. By utilising, comparing and combining the strengths of low error rates and high sequencing depth in Illumina RNA sequencing, and the full-length transcript sequencing in PacBio’s Iso-Seq, this annotation culminates in a merged transcriptome with functional annotation and an expression atlas. Evidence from small RNA of the same tissues sequenced using the Illumina platform revealed small RNAs that would have otherwise remained undetected. Our findings from a comparison between short-read and long-read reference transcriptomics contribute to a deeper understanding of these competing options. In this study, both technologies complemented each other. While short-read data was sufficient to annotate protein-coding genes, long-read data recovered more transcripts per gene and potentially further protein-coding genes that could not be annotated. With the ongoing improvement of base call quality in long-read sequencing, short-read transcriptome sequencing might become expendable. Together, the genome and transcriptome annotation of the tufted duck is an excellent resource for public omics databases and a foundation for downstream studies, e.g., regarding disease response. The dataset’s high quality for a non-model species allows for a much finer resolution of genetic differences and commonalities in closely related species, which is crucial while studying the reservoirs of zoonotic pathogens.

## Methods

### Sampling and dissection of tissues

Captive-bred, wild tufted ducks were kept at the animal breeding facility (Swedish National Veterinary Institute, Uppsala, Sweden). The ducks were obtained from Snavelhof breeding farm, Veeningen, the Netherlands, in May 2017. Tissue samples were obtained from five females and five males (12 months old) after euthanasia with an injection of 1 mL of pentobarbital (100 mg) in the wing vein. The following tissues were collected from the birds: brain, ileum, spleen, lung and gonads (ovary or testis). Tissues were immediately snap-frozen in liquid nitrogen and stored at −80°C until shipment on dry ice to the Roslin Institute, Edinburgh, UK. All animal experiments were carried out in strict accordance with a protocol legally approved by the regional board of the Uppsala animal ethics committee, Sweden (permission number 5.8.18-07998/2017). The animal experiments were conducted in biosafety level two (BSL-2) animal facilities at the Swedish National Veterinary Institute.

### Genomic DNA: library preparation, sequencing and assembly

To obtain both sex chromosomes, DNA was extracted from lung tissue of a female tufted duck. Library preparation and sequencing was conducted as in [21], using four types of sequencing data and the Vertebrate Genomes Project (VGP) assembly pipeline 1.6 (all details given in Rhie *et al*. [21]). The sequence data consisted of PacBio CLRs, 10X Genomics chromium linked-reads, Bionano Genomics optical maps created by direct labelling and staining (DLS), and chromatin conformation capture coupled with high throughput sequencing (Hi-C) (Arima Genomics, San Diego, CA, USA). In brief, PacBio reads were input to the diploid-aware long-read assembler FALCON and its haplotype-resolving tool FALCON-Unzip [66]. The resulting primary contigs were input to the Purge-Dups pipeline [67] to identify and remove remaining haplotigs in the primary set. In the next step, primary-purged contigs were subject to two rounds of scaffolding using the 10X long molecule linked-reads. Further, pre-assembled DLS Bionano cmaps were applied for further scaffolding and ordering using the Solve pipeline (Bionano Genomics, San Diego, CA, USA). The resulting scaffolds were then further scaffolded into chromosome-scale scaffolds using Hi-C data and the Salsa2 pipeline [68]. Finally, the primary assembly plus the Falcon-phased haplotigs were concatenated for three rounds of base call polishing: first with PacBio reads and Arrow software [69], and subsequently two rounds of polishing with 10X linked-reads and FreeBayes software [70]. The genome was decontaminated and manually curated as described in Howe *et al*. [71].

### Tissue preparation, DNA and RNA extraction

For disruption and homogenisation of tissues, snap-frozen samples were ground to a fine powder under liquid nitrogen using a mortar and pestle. Samples were transferred to 1.5 mL frozen tubes and kept on dry ice until further processing. Total RNA was obtained following a standard TRIzol protocol with DNase treatment and column purification. Small RNA was prepared according to the miRNeasy kit protocol 217004 (Qiagen, Venlo, Netherlands). Integrity and quality of the RNA were confirmed by electrophoresis on an Agilent 2200 Tapestation using appropriate screen tapes. The concentration was determined using the Nanodrop ND-1000 (Thermo Fisher Scientific, Waltham, MA, USA) (Tab. S1, Tab S2). For DNA extraction and sequencing, powdered lung tissue was sent to the Vertebrate Genomes Lab (VGL) at Rockefeller University, New York, USA.

### Illumina cDNA library preparation and sequencing

RNA was sent to Edinburgh Genomics, Edinburgh, UK, for library preparation and sequencing on an Illumina HiSeq 4000 platform with 2 x 150 bp paired-end reads using the TruSeq library preparation protocol (stranded). Median insert size was 137 bp - 148 bp (SD: 67 bp - 81 bp), yielding at least 290 M + 290 M reads per sample. Small RNA was also sent to Edinburgh Genomics for TruSeq small RNA library preparation and sequencing using a NovaSeq 6000 platform with 2 x 50 bp paired-end reads. Median insert size was 141 bp - 144 bp yielding at least 225 M + 225 M reads per sample.

### PacBio cDNA library preparation and sequencing

Two micrograms of total RNA from each sample in four parallel reactions were converted to cDNA using the Teloprime fulllength cDNA amplification kit (013, v1) according to the manufacturer’s instructions (Lexogen, Vienna, Austria). After end-point PCRs, all samples were tested for quality and quantity. The product size distribution was visualised using an Agilent 2200 Tapestation using D5000 screen tape. The library concentration was measured on a Qubit 3 (Thermo Fisher Scientific, Waltham, MA, USA) with high-sensitivity DNA reagents (Tab. S3). Technical replicates were pooled and selected for PacBio Iso-Seq assays. The samples were sequenced at Edinburgh Genomics using Sequel (version 2.1) chemistry.

### Gene/transcript annotation

Illumina raw RNA-Seq reads were quality checked and filtered using Fastqc (0.11.8) [72] and Trimmomatic (0.38) [73], respectively. Corrected reads were mapped to the genome using HISAT2 (2.2.0) [74, 75] and transcript models assembled using StringTie2 (2.1.1) [47]. The resulting annotation file was converted with tama_format_gtf_to_bed12_stringtie.py. In the remainder of this manuscript, all tools described as starting with “tama_” are part of the software suite Transcriptome Annotation by Modular Algorithms [76], except for tama_merge_report_parser.pl [77].

PacBio raw Iso-Seq reads were pre-processed using the IsoSeq3 pipeline to obtain full-length, non-chimeric reads (FLNC; first three steps in [78]; ccs 3.3.0, lima 1.8.0, refine 3.1.0). Afterwards, fasta sequences were extracted from bam files using Bamtools 2.5.1 [79] and poly-A tails were trimmed using tama_flnc_polya_cleanup.py (20191022). These FLNC were mapped to the reference genome with the splice site-aware mapper Minimap2 2.17-r974-dirty [80]. Redundant transcript models were collapsed with the capped flag (-x capped) using tama_collapse.py when coverage was at least 95% (-c 95). Additionally, 5’ threshold and 3’ threshold (tolerance in bp for grouping reads) were set to 100 (-a 100 -z 100).

### Functional annotation

Transcriptomes from all six tissues inferred by the short-read and long-read pipelines were merged based on similarity using tama_merge.py (options -a 100 -z 100 -d merge_dup) with different priorities for splice junctions and transcript end sites. Short-read inferred transcript models were given higher priority on splice junctions, whereas long-read inferred transcript models were given higher priority on transcript end sites. Nucleotide sequences based on coordinates in the merged transcriptome were extracted from the reference genome using Bedtools (2.29.0) [81]. The protein-coding potential was predicted with CPC2 (0.1) [82] based on the transcripts’ sequence features. Open reading frames (ORFs) were predicted and translated into amino acid sequences using tama_orf_seeker.py. Putative protein-coding sequences were identified in the UniProt/UniRef50 database (2019_10) [83] using Blastp (2.9.0) [84]. The results were filtered for top hits with tama_orf_blastp_parser.py, and a new annotation with coding sequence (CDS) regions was created using tama_cds_regions_bed_add.py.

### Tissue-specific expression analysis

In addition to merging annotations, tama_merge.py also creates a gene report containing each feature’s source (in this case: pipeline and tissue). This gene report was parsed with tama_merge_report_parser.pl [77] and filtered for genes found in UniRef50: Each feature was assigned a binary TRUE or FALSE label depending on the support of each of the twelve sources (two pipelines and six tissues). The result was loaded into Upsetr [85, 86] to visualise intersections of tissue-specific evidence for an expressed gene in each pipeline.

### Small RNA analyses

Illumina raw reads were quality checked with Fastqc (0.11.8) [72] and adapters removed with Cutadapt (2.10) [87]. Corrected reads were mapped to the reference genome using STAR (2.7.3a) [88] and assembled with Cufflinks (2.2.1) [89–92]. Nucleotide sequences were extracted from the reference genome at the assembled transcripts’ coordinates using Bedtools getfasta -split (2.29.2), and transcript lengths extracted from the output of Samtools faidx [93]. All plots were created using the package ggplot2 [94] in RStudio [86, 95].

The tool cmscan (1.1.3) from the software suite Infernal [96] was used to predict structural RNAs in the reference genome *(in silico)* and to annotate assembled small RNA transcripts *(in vitro)* based on Rfam [97, 98] covariance models (CM) downloaded from ftp://ftp.ebi.ac.uk/pub/databases/Rfam/14.2.

The output of cmscan (tblout) was converted to gff3 annotation files using tblout2gff3.pl [77]. Shared intervals between *in silico* and *in vitro* annotations were identified with the intersect option of Bedtools (2.29.2) [81].

## Supporting information

Supplementary data, Tufted Duck Annotation

## Abbreviations

AIV: Avian influenza A virus |
BSL: Biosafety level |
CLR: Continuous long reads |
DLS: Direct label and stain |
FLNC: Full-length, non-chimeric reads |
HPAI: Highly pathogenic avian influenza |
NGS: Next-generation sequencing |
ORF: Open reading frame |
SMRT: Single-molecule, real-time |
TAMA: Transcriptome annotation by modular algorithms |
VGL: Vertebrate Genomes Lab |
VGP: Vertebrate Genomes Project |
ZMW: Zero-mode waveguide

## Declerations

### Ethics approval and consent to participate

All animal experiments were carried out in strict accordance with a protocol legally approved by the regional board of the animal ethics committee, Sweden (permission number 5.8.18-07998/2017). The animal experiments were conducted in biosafety level two (BSL-2) animal facilities at the Swedish National Veterinary Institute (Uppsala, Sweden).

### Consent for publication

Not applicable

### Availability of source code and requirements

Perl and R scripts used in this study are available on GitLab at https://gitlab.com/rcmueller/tufted_duck_annotation under MIT license.

### Availability of data and materials

All RNA sequencing raw reads and data sets supporting the results of this article will be released upon acceptance.

### Funding

The authors gratefully acknowledge funding of a Short-Term Scientific Mission (STSM) through COST Action CA15112- Functional Annotation of Animal Genomes - European network (FAANG-Europe), and funding from the Ministry of Science, Research and the Arts Baden-Württemberg (MWK) during the Science Data Centre project BioDATEN.

## Author’s contributions

RCM: Conceptualisation, data curation, formal analysis, investigation, methodology, project administration, software, visualisation, writing–original draft, writing–review & editing | PE: Resources, writing–original draft | KH: Formal analysis, methodology, resources, validation, visualisation, writing–original draft | MUS: Formal analysis, methodology, validation, writing–original draft | RIK: Conceptualisation, methodology, software | KM: Methodology, writing–original draft | AW: Resources | OF: Formal analysis | BH: Formal analysis | JM: Formal analysis | WC: Formal analysis | JT: Formal analysis | JW: Formal analysis | JDJ: Resources | MMN: Resources | BO: Funding acquisition | EDJ: Project administration, resources | JS: Project administration, resources, supervision | LE: Formal analysis, investigation, visualisation | RHSK: Conceptualisation, Funding acquisition, project administration, resources, supervision

All authors read and approved the final manuscript.

## Acknowledgements

The authors acknowledge support by the local HPC resources through the core facility SCCKN at the University of Konstanz. The authors also acknowledge continued support by the Max Planck Insitute of Animal Behavior and its director Martin Wikelski.

