## Supplementary data, Tufted Duck Annotation for "A high-quality Genome and Comparison of Short versus Long Read Transcriptome of the Palaearctic duck *Aythya fuligula* (Tufted Duck)"

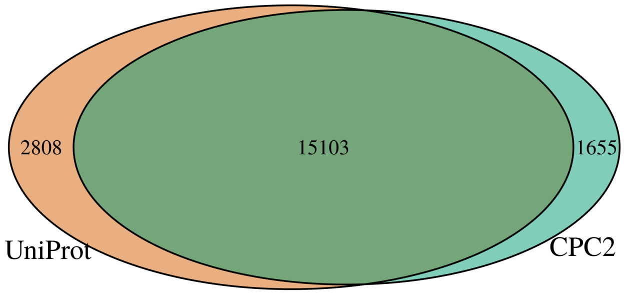


**Fig. S1.** Protein-coding potential calculated by CPC2 intersected with hits against the UniRef50 database (merged transcripts of all tissues and pipelines).

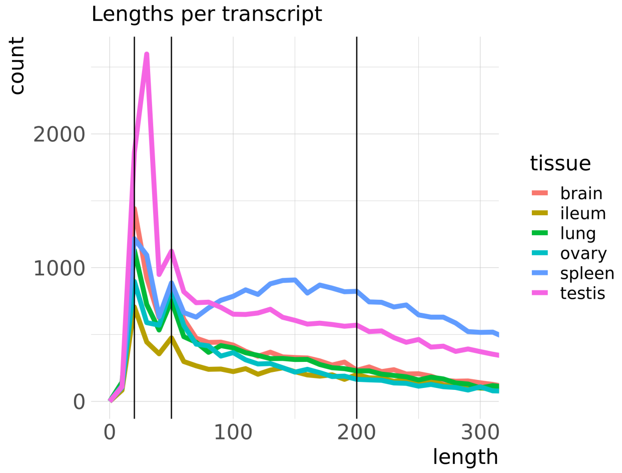


**Fig. S2.** Length distribution of small RNA transcripts follow decaying function with peaks at 20 bp (30 bp in testis) and 50 bp. Vertical bars at 20 bp, 50 bp and 200 bp were added.

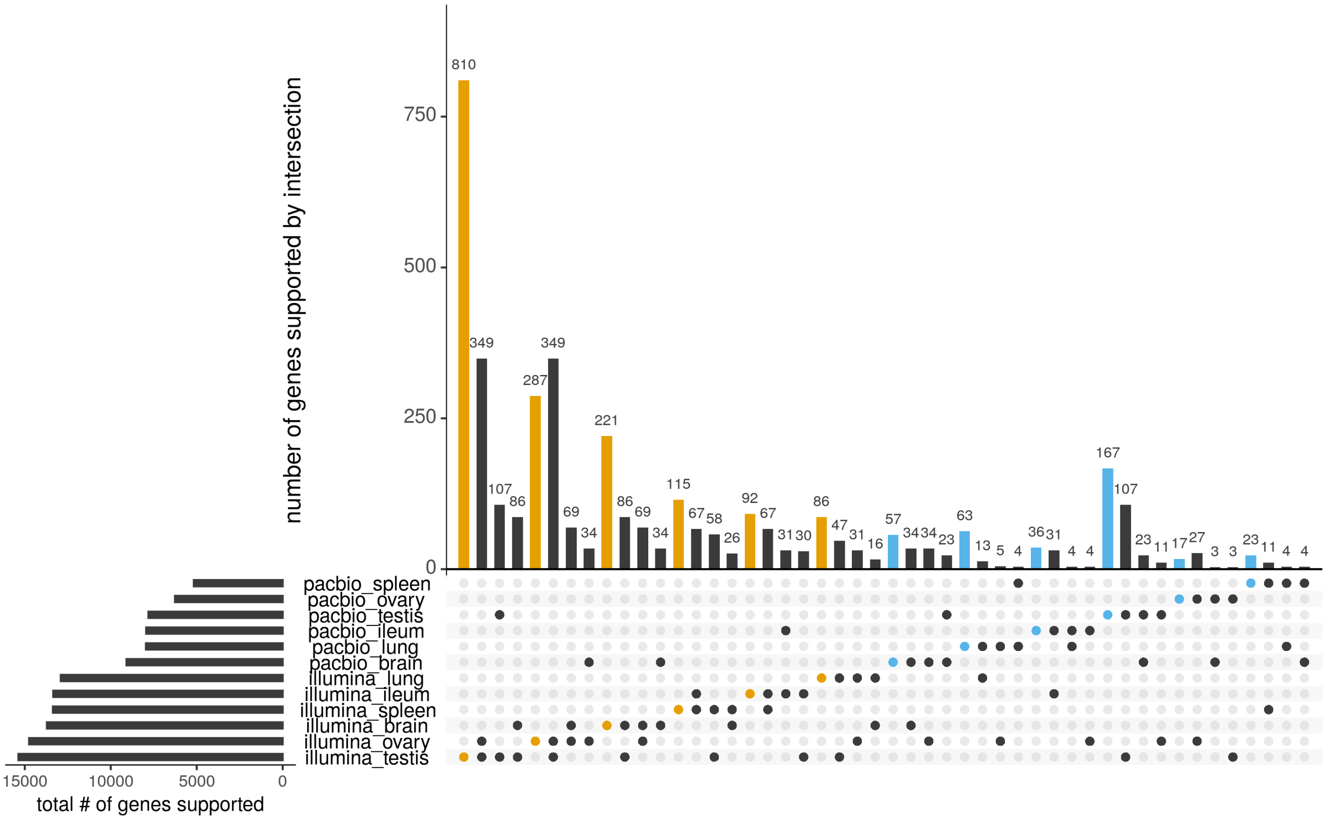


**Fig. S3.** Pipeline-tissue-specific expression of genes found in UniProt: Tissue-exclusive Illumina genes marked yellow, tissue-exclusive PacBio genes marked blue. Bottom left panel: total number of identified genes by pipeline and tissue ordered by the number of identified genes. Main panel: 12 grouped sets (pipeline-tissue) with exclusive genes plus top three intersections with other pipeline or tissue. The number of annotated genes was generally higher with Illumina transcriptome reconstruction than with that of PacBio. In both pipelines, testis expressed the highest number of genes exclusively, with 810 in Illumina data and 167 from PacBio.

**Tab. S1.** RNA extraction

| Sample | Weight [mg] | Concentration [ng/µL] | RIN |
|---|---|---|---|
| TD_brain_52037 | 123 | 202.7 | 9.2 |
| TD_ileum_52037 | 42 | 958.2 | 10.0 |
| TD_lung_52037 | 37 | 221.6 | 9.5 |
| TD_ovary_52037 | 55 | 1037.4 | 9.5 |
| TD_spleen_52037 | 53 | 2109.8 | 9.4 |
| TD_testis_54068 | 63 | 989.3 | 9.7 |

**Tab. S2.** small RNA extraction

| Sample | Weight [mg] | Concentration [ng/µL] | RIN |
|---|---|---|---|
| TD_brain_F52037 | 29 | 852.87 | 9.3 |
| TD_ileum_F52037 | 16 | 727.93 | 10.0 |
| TD_lung_F50325 | 15 | 402.63 | 9.7 |
| TD_ovary_F50325 | 12 | 368.79 | 9.4 |
| TD_spleen_F50325 | 5 | 823.70 | 9.5 |
| TD_testis_M54068 | 19 | 722.62 | 9.4 |

**Tab. S3.** cDNA libraries for Iso-Seq

| Sample | Concentration [ng/µL] |
|---|---|
| TD_brain_52037 | 46.2 |
| TD_ileum_52037 | 43.2 |
| TD_lung_52037 | 33.2 |
| TD_ovary_52037 | 41.0 |
| TD_spleen_52037 | 66.8 |
| TD_testis_54068 | 45.2 |
